# The Plasminogen-Apple-Nematode (PAN) domain suppresses JA/ET defense pathways in plants

**DOI:** 10.1101/2023.06.15.545202

**Authors:** Kuntal De, Debjani Pal, Carly M. Shanks, Timothy B. Yates, Kai Feng, Sara S. Jawdy, Md Mahmudul Hassan, Pradeep K. Prabhakar, Jeong-Yeh Yang, Digantkumar Chapla, Kelley W. Moremen, Breeanna Urbanowicz, Brad M Binder, Wellington Muchero

## Abstract

Suppression of immune response is a phenomenon that enables biological processes such as gamete fertilization, cell growth, cell proliferation, endophyte recruitment, parasitism, and pathogenesis. Here, we show for the first time that the Plasminogen-Apple-Nematode (PAN) domain present in G-type lectin receptor-like kinases is essential for immunosuppression in plants. Defense pathways involving jasmonic acid and ethylene are critical for plant immunity against microbes, necrotrophic pathogens, parasites, and insects. Using two *Salix purpurea* G-type lectin receptor kinases, we demonstrated that intact PAN domains suppress jasmonic acid and ethylene signaling in Arabidopsis and tobacco. Variants of the same receptors with mutated residues in this domain could trigger induction of both defense pathways. Assessment of signaling processes revealed significant differences between receptors with intact and mutated PAN domain in MAPK phosphorylation, global transcriptional reprogramming, induction of downstream signaling components, hormone biosynthesis and resistance to *Botrytis cinerea*. Further, we demonstrated that the domain is required for oligomerization, ubiquitination, and proteolytic degradation of these receptors. These processes were completely disrupted when conserved residues in the domain were mutated. Additionally, we have tested the hypothesis in recently characterized *Arabidopsis* mutant which has predicted PAN domain and negatively regulates plant immunity against root nematodes. *ern1.1* mutant complemented with mutated PAN shows triggered immune response with elevated WRKY33 expression, hyperphosphorylation of MAPK and resistant to necrotrophic fungus *Botrytis cinerea*. Collectively, our results suggest that ubiquitination and proteolytic degradation mediated by the PAN domain plays a role in receptor turn-over to suppress jasmonic acid and ethylene defense signaling in plants.

## Introduction

The effectiveness of plant immunity largely depends on its fine-tuning between growth and defense mechanism. Due to the lack of “mobile immune cells,” plants have evolved highly complex and regulated cell surface surveillance mechanisms, primarily consisting of different receptor-like kinases (RLK) and receptor-like proteins (RLP) together known as pattern recognition receptors (PRR). In general, a plant RLK contains an extracellular ligand recognition domain followed by a transmembrane domain and an intracellular kinase domain. Based on the structure of the extracellular domain, the RLKs are divided into subclasses. One of the most crucial ones for plant immunity is the Lectin-RLKs with a distinct N-terminal Lectin domain. Following activation, these kinases turn on a series of downstream signaling pathways working together with numerous cellular processes to establish immunogenic responses. However, emerging evidence indicates that the acquired immunity to protect themselves from the attacking pathogens is far more complex as plants need to balance both beneficial and pathogenic invaders as the former provides essential nutrients and is vital for agriculture outcomes. In contrast, the latter is solely responsible for different diseases and crop destructions. It is of great interest to know whether the cell surface recognition components determine the pattern of the immunity responses or whether microbes have evolved a mechanism to overcome it. Suppression of immune response is a well-known phenomenon that is involved in biological processes including gamete fertilization, cell growth and proliferation, recruitment of symbionts, parasitism, and pathogenesis. As such understanding the molecular basis of this phenomenon is of crucial importance. In our recent study, we reported the occurrence of the plasminogen-apple-nematode (PAN) domain in more the 28,000 proteins across 2,496 organisms representing 959 genera. A GO enrichment analysis in that study revealed that these proteins were highly enriched in terms related to immune responses including cell recognition, cell communication, proteolysis, pollen pistil recognition, reproduction, responses to stimulus and response to stress (Pal, De et al. 2022). Among the >28,000 PAN domain-containing proteins, the only plant protein family represented were G-type lectin receptor-like kinases (G-type LecRLKs). G-type LecRLKs are one of the three subtypes of LecRLKs with an ectodomain resembling like Galanthus nivalis agglutinin (GNA) mannose binding motif. G-type LecRLKs also have the highest copy numbers across plant genomes compared to their C- and L-type LecRLK counterparts. One of the distinguished features for G-type LecRLks is to have the plasminogen-apple-nematode (PAN) domain. The term PAN domain came from the work of Tordai et al. back in 1999 as it was first discovered in plasminogen/hepatocyte growth factor family (HGF) and predicted to be serve as a site for protein-protein or protein-carbohydrate interaction. We have shown direct evidence that the PAN domain of HGF acts as a binding site for its receptor, c-MET and regulate the downstream signaling cascade including ERK phosphorylation (Pal, De et al. 2022).

Interestingly, members of the G-type LecRLK family are increasingly implicated in enhancing microbial colonization or parasitic infection in plants. For example, a study showed that constitutive expression of the *Populus trichocarpa* PtG-type LecRLK1 in Arabidopsis resulted in enhanced colonization by the ectomycorrhizal fungal symbiont Laccaria bicolor, even though Arabidopsis is not a natural host of this fungus (Labbé, Muchero et al. 2019, Qiao, Yates et al. 2021). In a follow-up study, the same *Pt*G-type LecRLK1 was expressed in switchgrass resulting in the successful formation of functional mycorrhizae (Qiao, Yates et al. 2021). The latter result is especially noteworthy since grasses are not known to form mycorrhizae with ectomycorrhizal fungi. In another study, a G-type LecRLK was reported to function as a susceptibility factor to the fungal pathogen *Sphaerulina musiva* in *P. trichocarpa* (Muchero, Sondreli et al. 2018). Besides fungal symbionts and pathogens, a G-type LecRLK was also shown to mediate infection of *Arabidopsis* by parasitic root-knot nematodes. Loss-of-function mutants of this receptor resulted in enhanced resistance to nematode infection, supporting the role of G-type LecRLKs as negative regulators of immune responses (Zhou, Godinez-Vidal et al. 2021). Despite these observed instances of enhanced susceptibility facilitated by G-type LecRLKs, the mechanism by which these receptors function to repress defense signaling and facilitate infection remains entirely unknown. JA is a key regulator for the activation of plant defense pathway usually synthesized in plants during the attack by invading pathogens. The activation of JA and ET pathway leads to the upregulation of pathogen-responsive genes in plants, ultimately counteracting disease progression. This study, for the first time, demonstrates that G-type LecRLKs suppress jasmonic acid (JA) and ethylene (ET) signaling pathways via the PAN domain. We further defined that intact PAN domain attenuates plant immunity by inhibiting MAP kinase phosphorylation and reactive oxygen species (ROS) production. Additionally, we find that conserved PAN domain is required for the ubiquitin mediated proteasomal degradation of the G-type LecRLK. We have also tested the function of PAN domain in *Arabidopsis* G-LecRLK receptor protein, ERN1 which shows that mutation in the domain elevates immune response in plants. Together our data provide a model whereby PAN domain acts as an upstream modulator that tunes the amplitude of JA/ET dependent plant immunity by controlling G-type LecRLK stability.

### Conserved PAN domain residues modulate WRKY33 expression

In this study, we characterized two *Salix purpurea* G-type LecRLKs, *Sp*G-type LecRLK-1 (SapurV1A.0918s0020) and *Sp*G-type LecRLK-2 (SapurV1A.003s0270) which, unlike their previously published counterparts, are fully capable of initiating WRKY33-mediated immune responses. We performed protein domain prediction on *Sp*G-type LecRLK-1 and *Sp*G-type LecRLK-2 using the InterPro database. This analysis suggested that both receptors were missing the PAN domain, which is uncharacteristic for this protein family. We then aligned *Sp*G-type LecRLK-1 and *Sp*G-type LecRLK-2 with characterized G-type LecRLKs from *Populus* and Arabidopsis. The alignment revealed that both receptors had sequences consistent with the PAN domain but had acquired mutations at the same seven conserved residues which resulted in statistically insignificant prediction of the domain. Specifically, *Sp*G-type LecRLK-1 had K19V, D27G, N50D, C51T, L69T, W71K and L75F while *Sp*G-type LecRLK-2 had K19T, D27G, N50D, C51P, L69R, W71K and L75Y mutations within their respective PAN domains (Figure 1A). Unlike typical G-type LecRLKs which do not elicit immune signaling, these two receptors surprisingly induced expression of the signature defense-related transcription factor, WRKY33, upon expression in Arabidopsis (Figure S1A & B) which is the signature gene for Jasmonic acid (JA)/ Ethylene (ET) pathway. However, we did not see any significant changes for NPR1 gene, which is a signature gene for Salicylic acid (SA) pathway (Figure S1A). As a comparator, we selected another *Salix* G-type LecRLK, SapurV1A.1175s0020, which has a predicted intact PAN domain. SapurV1A.1175s0020 could not induce WRKY33 expression upon overexpression in Arabidopsis (Figure S1B) which suggests that mutated version of PAN domain could be responsible for immune activation in plants. Thus, our observation suggested that the PAN domain may play a central role in repression of immune responses in plants.

**Figure 1:**
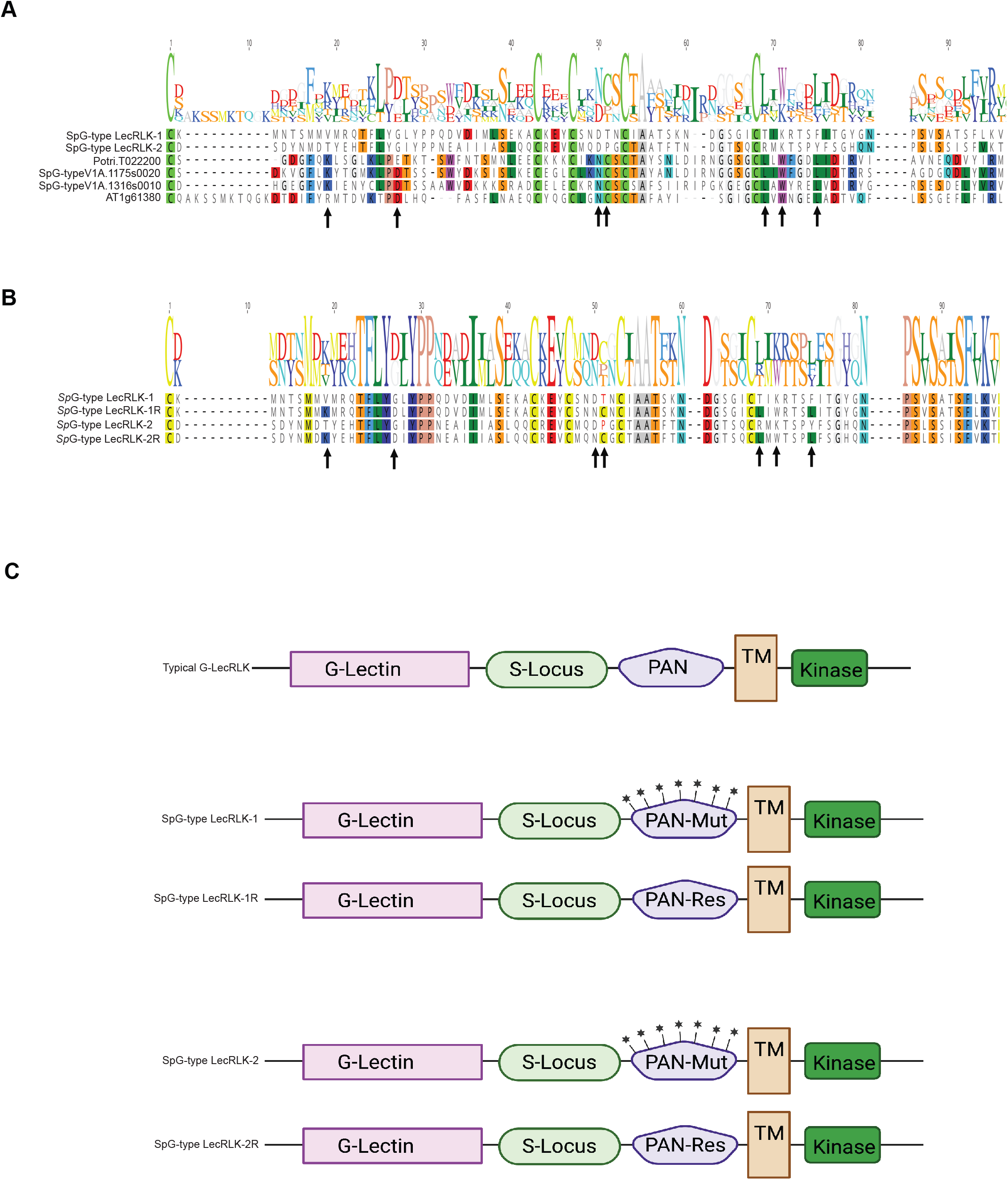
GLecRLK PAN domain is comprised of conserved amino acid residues and is critical for immunosuppression in plants. A. Multiple sequence alignment within the GLecRK PAN domains and naturally occurring variation in non-immunosuppressive Salix orthologs SapurV1A.0918s0020 (SpG-type LecRLK-1) and SapurV1A.0037s0270.1 (SpG-type LecRLK-2). B. Multiple sequence alignment with the G-LecRK PAN domains and naturally occurring variation in non-immunosuppressive Salix orthologs SapurV1A.0918s0020 (SpG-type LecRLK-1) and SapurV1A.0037s0270.1 (SpG-type LecRLK-2) with their respective restored version. C. Schematic representation of typical G-type LecRLKs, G-type LecRLKs with PAN domain mutations (*Sp*G-type LecRLK-1 & *Sp*G-type LecRLK-2) and G-type LecRLKs with restored PAN domains (*Sp*G-type LecRLK-1R & *Sp*G-type LecRLK-2R).

To test the hypothesis that mutated residues in the PAN domain are essential for negative defense regulation, we created variants of the two receptors with mutated amino acids restored to conserved residues and designated them *Sp*G-type LecRLK-1R and *Sp*G-type LecRLK-2R (Figure 1B). *Arabidopsis thaliana* transgenic plants constitutively expressing the four protein variants were created for further characterization (Figure S1C). The transgenic variants do not show any visible morphological difference (Figure S1D). Figure 1C represents a schematic of typical G-type LecRLKs, G-type LecRLKs with PAN domain mutations (*Sp*G-type LecRLK-1 & *Sp*G-type LecRLK-2) and G-type LecRLKs with restored PAN domains (*Sp*G-type LecRLK-1R & *Sp*G-type LecRLK-2R). We also created tobacco (*Nicotiana benthamiana*) expressing *Sp*G-type LecRLK-1 and its restored version SpG-type LecRLK-1R (Figure S1E). qPCR analysis and transcriptional profiling using RNA-seq of transgenic plants confirmed our observations that *Sp*G-type LecRLK-1 and *Sp*G-type LecRLK-2 could significantly induce WRKY33 expression. In contrast, the two restored variants, *Sp*G-type LecRLK-1R and *Sp*G-type LecRLK-2R, lost their ability to induce WRKY33 expression (Figure 2A and Figure 2B) and (Figure S1C and Figure S1E).

**Figure 2:**
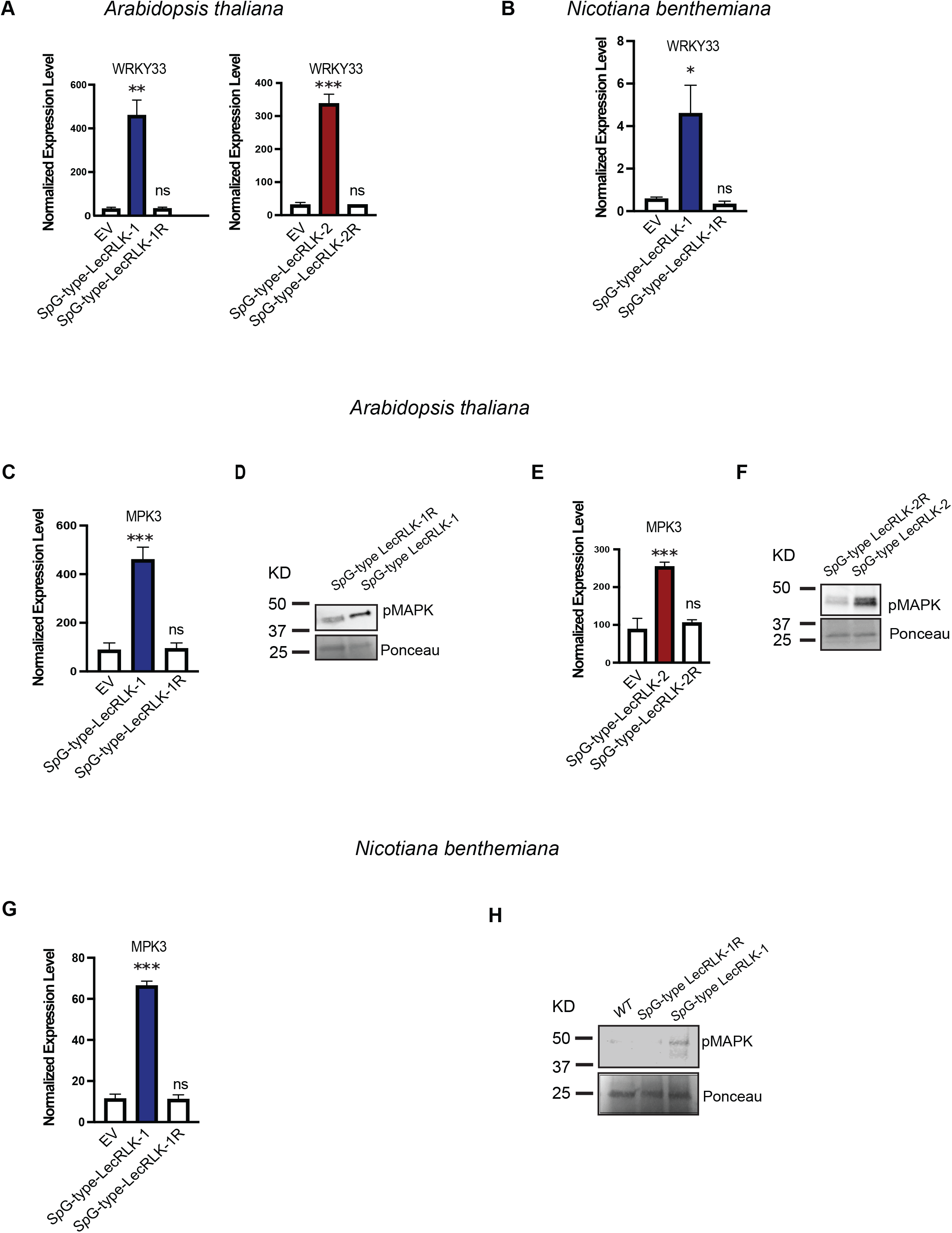
G-type RLKs suppresses WRKY33 transcription. A. Transcriptome data from Arabidopsis transgenic plants shows differential expression of WRKY33 in transgenic Arabidopsis plants. Results are shown as mean ± SE of four different independent experiments. Asterisks represents significant differences between transgenic plants and empty vector control (***P<0.0005; **P<.005; *P<0.05 Student T-test). B. Transcriptome data from Nicotiana transgenic plants shows differential expression of WRKY33. Results are shown as mean ± SE of three different independent experiments. Asterisks represents significant differences between transgenic plants and wild type control (***P<0.0005; **P<.005; *P<0.05 Student T-test). C, and E. Transcriptome data from Arabidopsis transgenic plants shows differential expression of MPK3. Results are shown as mean ± SE of four different independent experiments. Asterisks represents significant differences between transgenic plants and empty vector control (***P<0.0005;**P<.005; *P<0.05 Student T-test). D and F. Western blot analysis demonstrated MAPK is hyperphosphorylated in transgenic SpGtype LecRLK-1 and SpG-type LecRLK-2 Arabidopsis compared to SpG-type LecRLK-1R and SpG-type LecRLK-2R Arabidopsis plants. Representative image of 3 independent experiments which shows similar results. G. Transcriptome data from Nicotiana transgenic plants shows MPK3 is significantly upregulated in SpG-type LecRLK-1. H. Western blot analysis demonstrated MAPK is hyperphosphorylated in transgenic SpG-type LecRLK-1 compared to SpG-type LecRLK-1R Nicotiana plants. Representative image of 2 independent experiments which shows similar results.

To investigate the subcellular localization of *Sp*G-typeRLK-1, *Sp*G-typeRLK-2 and its restored version, we generated constructs expressing a translational fusion of *Sp*G-typeRLK-1 with GFP and *Sp*G-typeRLK-1R with GFP under the control of strong cauliflower mosaic virus (CaMV) 35S promoter. We then tested the transient expression in tobacco using *Agrobacterium tumefaciens* infiltration approach. In this experiment, both *Sp*G-typeRLK-1 (Figure S2B) and *Sp*G-typeRLK-1R (Figure S2C) appeared to be localized to the PM of leaf cells, although GFP alone appeared to be present in both cytoplasm and nucleus (Figure S2A and Figure S2D). Similarly, *Sp*G-typeRLK-2 GFP (Figure S2E) and *Sp*G-typeRLK-2R GFP (Figure S2F) showed similar pattern of localization to the PM of the leaf cells. From this experiment, we conclude that there is no difference in localization patterns between the mutated and restored versions of PAN domain.

### PAN domain controls MAPK expression and phosphorylation in plants

We sought to assess the status of the JA/ET signaling cascades downstream of the four receptor variants. Firstly, we evaluated expression of mitogen activated protein kinases (MAPKs) which are phosphorylation targets of the kinase domain of membrane-bound receptors. Two well-known MAPKs, MPK3 and MPK6, have been implicated in defense against pathogens (Asai, Tena et al. 2002, Ichimura, Casais et al. 2006, Qiu, Fiil et al. 2008). Transcriptome data from Arabidopsis transgenic plants revealed that expression of MPK3 was significantly higher in transgenic *Sp*G-type LecRLK-1(Figure 2C) and *Sp*G-type LecRLK-2 (Figure 2E) compared to wildtype or the transgenics expressing restored variants. Similar trend we determined from tobacco transgenic *Sp*G-type LecRLK-1 plants (Figure 2G). Additionally, western blot analysis demonstrated that MAPK was hyperphosphorylated in Arabidopsis expressing *Sp*G-type LecRLK-1 (Figure 2D) and *Sp*G-type LecRLK-2 (Figure 2F) as well as in tobacco expressing *SpG*-type LecRLK-1(Figure 2H). However, notably reduced phosphorylation was observed in transgenics expressing restored variant (Figure 2D, 2F, and 2H). We conclude that intact PAN domain suppresses pMAPK and is necessary to suppress immune activation.

### PAN domain specifically suppresses JA and ET pathway

Further, we examined expression levels of downstream genes that are known to mediate to JA and ET defense pathways. LOX3, LOX4, OPR3, JAZ1, JAZ7, AOC3, and PDF1.3 function in the JA pathway (Acosta and Farmer 2010), while ERF13, ERF1 and ACS6 function in the ET defense pathway (Dubois, Van den Broeck et al. 2018). In tobacco, we assayed expression levels of LOX3, OPR3, AOC3, JAZ1 and the ET-related transcription factor ERF1. All these genes exhibited the same expression pattern and were highly upregulated in *Sp*G-type LecRLK-1- and *Sp*G-type LecRLK-2-expressing Arabidopsis and tobacco transgenics but showing almost wild-type levels in Arabidopsis and tobacco expressing the restored variants, *Sp*G-type LecRLK-1R and *Sp*G-type LecRLK-2R (Figure 3A and Figure 3B).

**Figure 3:**
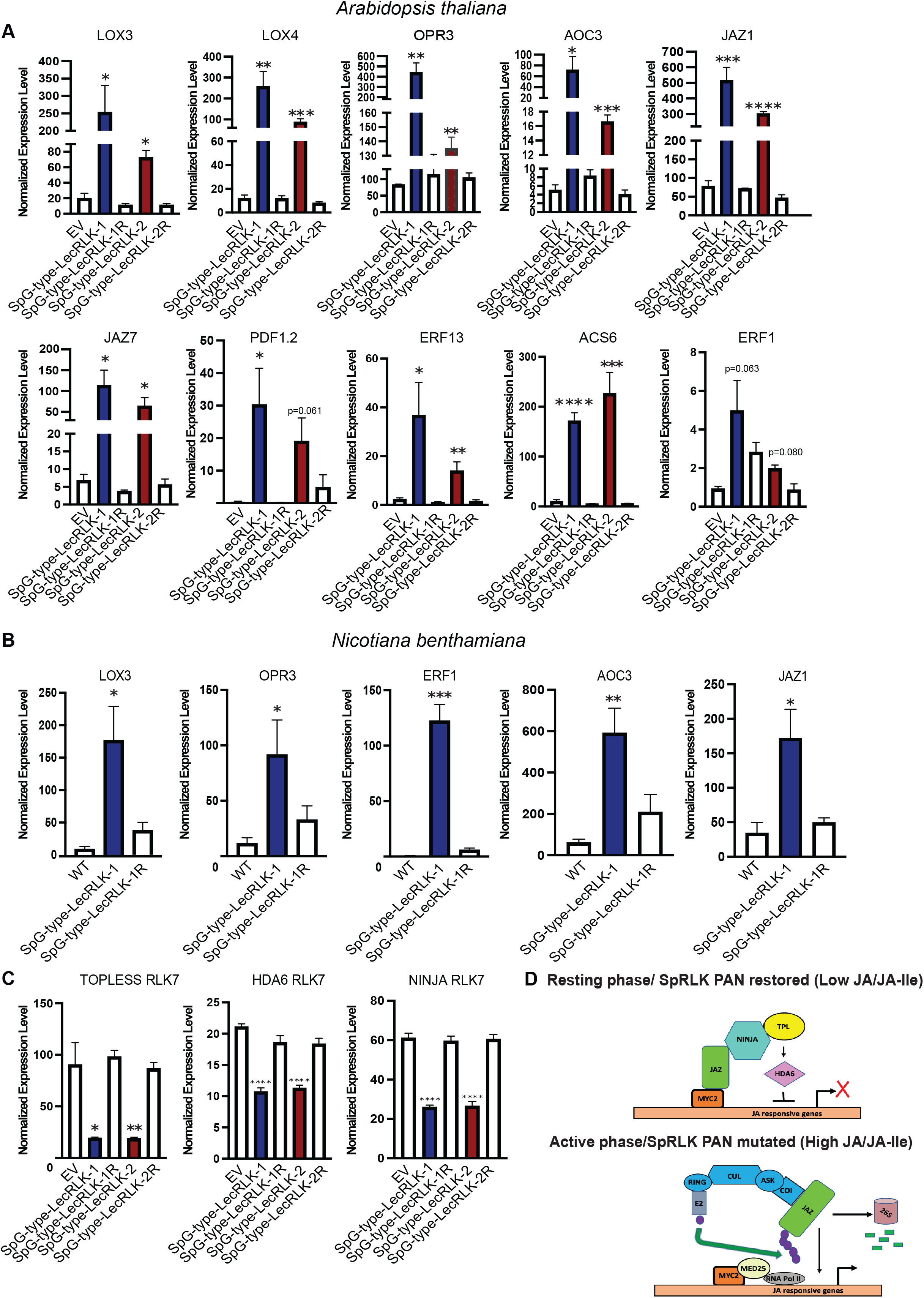
PAN domain specifically suppresses JA and ET pathway. A. Differential expression of genes involved in JA pathway and in ET pathway in Arabidopsis. Results are shown as mean ± SE (n=4 biological replicates). Asterisks represents significant differences between transgenic plants and empty vector control (***P<0.0005; **P<.005; *P<0.05 One-way Anova, Sidak method). B. Differential expression of genes involved in JA pathway and in ET pathway in Tobacco. Results are shown as mean ± SE (n=3 biological replicates). C. Reduced expression of negative regulators in mutated PAN transgenic version of SpG-type LecRLK-1 and SpG-type LecRLK-2. Results are shown as mean ± SE (n=4 biological replicates) (***P<0.0005; **P<.005; *P<0.05 One-way Anova, Sidak method). D. Mode of action showing mutating PAN domain repress TOPLESS, HDA6 and NINJA which ultimately triggers JA pathway.

We next extended our analysis of the effect of PAN domain on the transcriptional response of three well-known repressors, NINJA, HDA6 and TOPLESS, which function as negative regulators of the JA pathway. In contrast to JA and ET pathways gene above, all three repressors were significantly downregulated in *Sp*G-type LecRLK-1 and *Sp*G-type LecRLK-2 Arabidopsis transgenics while they were significantly higher in wildtype and transgenic plants expressing both restored variants (Figure 3C). These results confirmed that indeed the restored variants attenuated JA signaling compared to the variants with mutated PAN domains (Figure 3C). We also evaluated expression levels of U-box proteins PUB22 and PUB23 which are known for their role in regulating amplitude of receptor-triggered immune responses at the later stages of defense signaling cascades (Trujillo, Ichimura et al. 2008). Expression of both *Sp*G-type LecRLK-1- and *Sp*G-type LecRLK-2 resulted in increased induction of PUB22 and PUB23 (Figure S3A). In contrast, restored variants did not induce their expression confirming our claim that restored variants suppress defense signaling resulting in no observable immune responses (Figure S3A). JA and ET have also been shown to function antagonistically to abscisic acid (ABA) signaling (Anderson, Badruzsaufari et al. 2004). We therefore evaluated expression levels of OST1, a serine threonine protein kinase as well as VSP1 and VSP2, all known positive regulators of the ABA signaling pathway (Vos, Verhage et al. 2013). All three showed reduced expression in *Sp*G-type LecRLK-1 and *Sp*G-type LecRLK-2 transgenic lines while their expression was remarkably higher in wildtype and transgenics expressing restored variants *Sp*G-type LecRLK-1R and *Sp*G-type LecRLK-2R (Figure S3B).

Global differential expression analysis showed that 6,931 genes were differentially expressed in *Sp*G-type LecRLK-1 when compared to empty vector control (Figure S4A). However, we observed a significantly reduced number of genes (1,515) that were differentially expressed between the restored variant, *Sp*G-type LecRLK-1R compared to empty vector control (Figure S4A). Similarly, for *Sp*G-type LecRLK-2, 6,942 genes were differentially expressed compared to controls whereas 1,078 genes were differentially expressed in the restored variant *Sp*G-type LecRLK-2R compared to empty vector control (Figure S5A). GO enrichment analysis revealed that terms related to response to wounding, response to chitin, and regulation of jasmonic acid were enriched among differentially expressed genes, as indicated by green bullets (Figure S5B&C). These results support our hypothesis that the PAN domain plays a crucial role in repression of immune response in plants.

To provide definitive confirmation of differences in signaling between *Sp*G-type LecRLKs with mutated PAN domain amino acid residues and their restored variants, we performed phytohormone analysis and measured JA, jasmonoyl-isoleucine (JA-IIe), (SA) and (ABA) levels in Arabidopsis. Consistent with results of the transcriptome studies, JA and JA-IIe hormone levels were significantly higher in *Sp*G-type LecRLK-1 and *Sp*G-type LecRLK-2 transgenic Arabidopsis while the restored variants had empty vector control levels (Figure 4A and Figure 4B). SA and ABA exhibited negligible differences across empty vector control and all transgenics expressing the four variants (Figure 4C and Figure 4D). Several research studies also emphasized that the Jasmonic acid hormonal pathway and Ethylene pathway synergistically regulate plant immunity (Lorenzo, Piqueras et al. 2003)(Penninckx et al.1996, Dong 1998). We were also able to see the upregulation of ET-responsive genes which includes ERF13, and ERF1 in the *Sp*G-type LecRLK-1 and *Sp*G-type LecRLK-2 transgenic *Arabidopsis* lines (Figure 4E). We were interested in measuring the ethylene levels on these variants to confirm whether the change in the transcript level of the ET pathway produces the target phytohormone, ethylene. Our assay demonstrated that the level of ethylene phytohormone is significantly higher in the *Sp*G-type LecRLK-1 and *Sp*G-type LecRLK-2 transgenic *Arabidopsis* lines (Figure 4E). In contrast, the restored variants’ level changes are insignificant compared to the empty vector control (Figure 4E).

**Figure 4:**
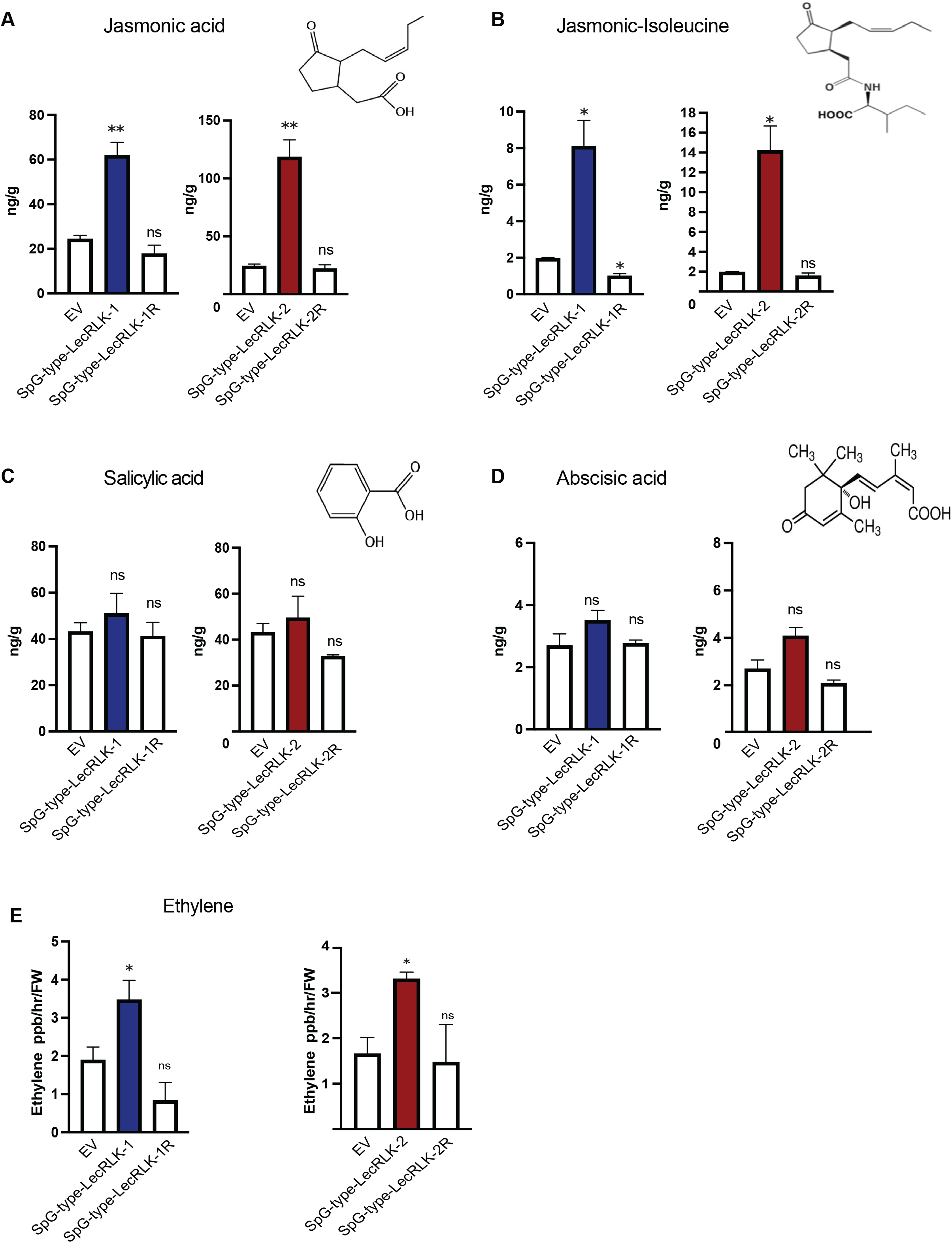
Functional PAN domain regulates the phytohormone level in plants. A. and B. Hormone analysis determines jasmonic acid and jasmonoyl-isoleucine (JA-IIe) hormone levels were significantly higher in SpG-type LecRLK-1 and SpG-type LecRLK-2 transgenic Arabidopsis while the restored variants had control levels. C and D. Salicylic acid and abscisic acid exhibited negligible changes in four variants. Results are shown as mean ± SE (n=3 biological replicates). Asterisks represents significant differences between transgenic plants and empty vector control (***P<0.0005; **P<.005; *P<0.05 Student T-test). E. Ethylene phytohormone analysis demonstarted significantly higher levels in SpG-type LecRLK-1 and SpG-type LecRLK-2 transgenic Arabidopsis.

Taken together the qPCR, RNA-seq, MAPK phosphorylation and hormone analyses described above demonstrate that minimal mutations that accumulated in the PAN domains of *Sp*G-type LecRLK-1 and *Sp*G-type LecRLK-2 were sufficient to interfere with their negative regulatory activity and restore JA and ET immune responses. Restoration of these mutations to their conserved amino acids reconstituted negative regulation of JA and ET immune signaling, suggesting an essential role of the PAN domain in this phenomenon.

### PAN domain modulates ROS response and immune suppression to *Botrytis cineara*

Given the cumulative observations above, we sought to establish functional consequences on defense against the pathogen, *Botrytis cinerea*. The receptor like cytoplasmic kinase *Botrytis* induced kinase 1 (BIK1) serves as a central regulator during PAMP-triggered immunity in plants (Jiang, Han et al. 2019). Phosphorylation of BIK1 induces its stability and activation, which in turn activates respiratory burst oxidase homolog protein D (RBOHD) (Jiang, Han et al. 2019). Activation of RBOHD releases reactive oxygen species (ROS) which is essential for plant immunity. Transcript levels of BIK1 and RBOHD were higher in both *Sp*G-type LecRLK-1 and *Sp*G-type LecRLK-2 transgenics compared to wildtype (Figure 5A and Figure 5B). As shown in Figure 5C and Figure 5D), we measured the oxidative burst triggered by the perception of flg22 in a luminol dependent assay. This analysis showed that *Sp*G-type LecRLK-1 and *Sp*G-type LecRLK-2 transgenic lines had distinct enhancement of the oxidative burst compared to wildtype or transgenics expressing restored variants (Figure 5C and 5D). Additionally, we demonstrated activation of ROS by analyzing leaves of *Sp*G-type LecRLK-1 transgenic lines using the dye H2DCFDA under confocal microscopy (Figure 5G). As our mutant version show higher expression of WRKY33, and ROS production, we were interested to evaluate whether the transgenic plants show any resistance against necrotrophic fungus, *Botrytis cinerea*. It is well known that transcription factor WRKY33 is essential for defense toward the fungus Botrytis cinerea(Birkenbihl, Diezel et al. 2012). In fact, Ethylene and jasmonic acid (ET/ JA) signaling are critical for plant immunity against necrotrophs (Glazebrook 2005, Bari and Jones 2009). Another study demonstrates that, transgenic *Arabidopsis* lines overexpressing *ethylene* responsive genes and signature genes in JA pat*hway* confer resistance to *B. cinerea* (Kazan and Manners 2013). Our pathogen assay determines, *Arabidopsis* transgenics expressing *Sp*G-type LecRLK-1 showed enhanced resistance against *Botrytis* compared to wildtype or its restored variant. Wildtype and *Sp*G-type LecRLK-1R transgenic leaves developed more necrosis compared to *Sp*G-type LecRLK-1 as depicted in (Figure 5E and Figure 5F). This pathogen assay conclusively demonstrated that JA and ET defense pathways were activated in *Sp*G-type LecRLKs with mutated PAN domains but were repressed upon restoration of PAN domain amino acids to conserved residues which is further depicted by a proposed model shown in (Figure 5H). Therefore, we conclude that the PAN domain underlies the observed negative regulation of JA and ET signaling by G-type LecRLKs in plants.

**Figure 5:**
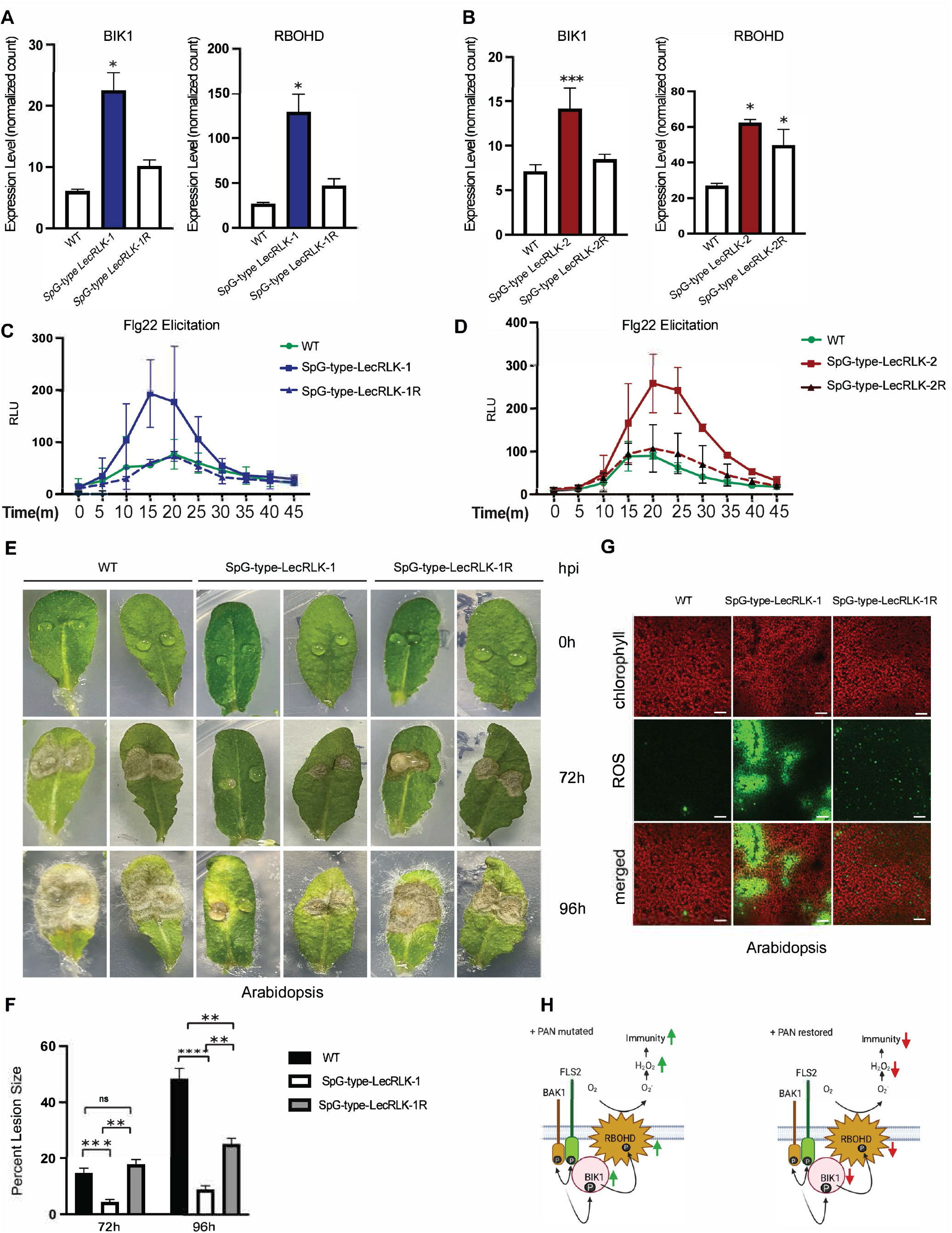
PAN domain modulates ROS response and immune suppression to Botrytis. (A and B). Expression level of two marker genes which includes Botrytis induced kinase 1 (BIK1) and RBOHD. Bars represent the standard error of the mean for four biological replicates. Statistical difference was determined by two-tailed student’s T test against Col-0 (***P<0.0005; **P<.005; *P<0.05 Student T-test). C and D. Flg22 elicitation assay: Oxidative burst triggered by 500nM flg22 in wild type Col-0, SpG-type LecRLK-1 and SpG-type LecRLK-1R transgenic plants measured in relative light units (RLU). Data represents standard deviation of the mean from three independent experiments from 9 biological replicates. D. Oxidative burst triggered by 500nM flg22 in wild type Col-0, SpGtype LecRLK-2 and SpG-type LecRLK-2R transgenic plants measured in relative light units (RLU). Data represents standard deviation of the mean from three independent experiments from 9 biological replicates. E. Botrytis infection assay: Botrytis cinerea inoculation of Col-0, SpG-type LecRLK-1 and SpGtype LecRLK-1R transgenic Arabidopsis plants. F. Lesion area (percentage of total leaf area) in 72 hours post inoculation and 96 hours post inoculation are represented in bar graphs. Value represents mean ± SE from at least n=20 from each group. Asterisks represents significant differences (***P<0.0005; **P<.005; *P<0.05 One-way Anova, Sidak method). G. Reactive oxygen species detection by confocal laser scanning microscopy in Arabidopsis leaves. Merged two-color confocal images of the green and the red channel show ROS detected by carboxy-H2DCFDA probe (green) and autofluorescence of chloroplast (red). The experiment is the representative image of 3 biological replicates. Size Bar:10um. H. Working model showing mutated PAN triggers immunity (left) and restored version dampens immunity (right).

### PAN domain self-interacts and modulates oligomerization

To provide mechanistic evidence for the basis of the observed differences between mutated *Sp*G-type LecRLKs and their restored variants, we performed biochemical experiments to establish any difference in protein behavior. Previously it has been demonstrated that the PAN domain mediated S-locus receptor dimerization in the absence of a ligand (Naithani, Chookajorn et al. 2007). Oligomerization of receptor-like kinases has been demonstrated to be essential for the activation of receptor kinases in plants and animals (Lemmon and Schlessinger 1994, Williams, Wilson et al. 1997). We therefore tested if there were any changes in the oligomeric state of mutated and restored forms of the two receptors. In this experiment, only the PAN domains were evaluated instead of full-length proteins.

Size exclusion chromatography indicates *Sp*G-type LecRLK-1PAN is present as a dimer or trimer/tetramer (Figure S6A). *Sp*G-type LecRLK-1RPAN expression was quite poor relative to its counterpart, and most of the protein aggregated, which is likely the reason for the low levels of secretion. The oligomeric profile of *Sp*G-type LecRLK-2PAN indicated that the protein was present in the solution is roughly split between monomeric (51.8%) and a higher-order complex that is likely a trimer or tetramer (48.2%) (Figure S6B). In contrast, only 26.4% of LecRLK-2RPAN was monomeric, and most of the protein formed higher-order complexes (Figure S6B), confirming the hypothesis that an intact PAN domain promotes receptor oligomerization. Additionally, to confirm further that *Sp*G-type LecRLK-1R promotes oligomerization we also expressed PAN domains of both *Sp*G-type LecRLK-1 and *Sp*G-type LecRLK-1R with different tags, GFP and FLAG in HEK293T cells. Co-immunoprecipitation results showed that PAN domains from both proteins were able to self-interact and the degree of interaction in the restored variants was higher compared to mutated forms (Figure S7A, S7B). These data also indicated that *Sp*G-type LecRLK-1 and *Sp*G-type LecRLK-2 have fewer oligomeric species compared to the restored variants.

### PAN domain conserved residues are critical for G-typeLecRLK stability

We were interested to evaluate the stability of the mutated and restored forms of PAN domain containing G-LecRLKs. To determine the mechanism, we tested the stability of the mutated and restored forms of both receptors using the cycloheximide chase assay in Arabidopsis protoplasts. Both *Sp*G-type LecRLK-1 and *Sp*G-type LecRLK-2 exhibited significantly extended stability over time compared to the restored variants *Sp*G-type LecRLK-1R, *Sp*G-type LecRLK-2R and empty vector control. (Figure 6A and Figure 6B).

**Figure 6:**
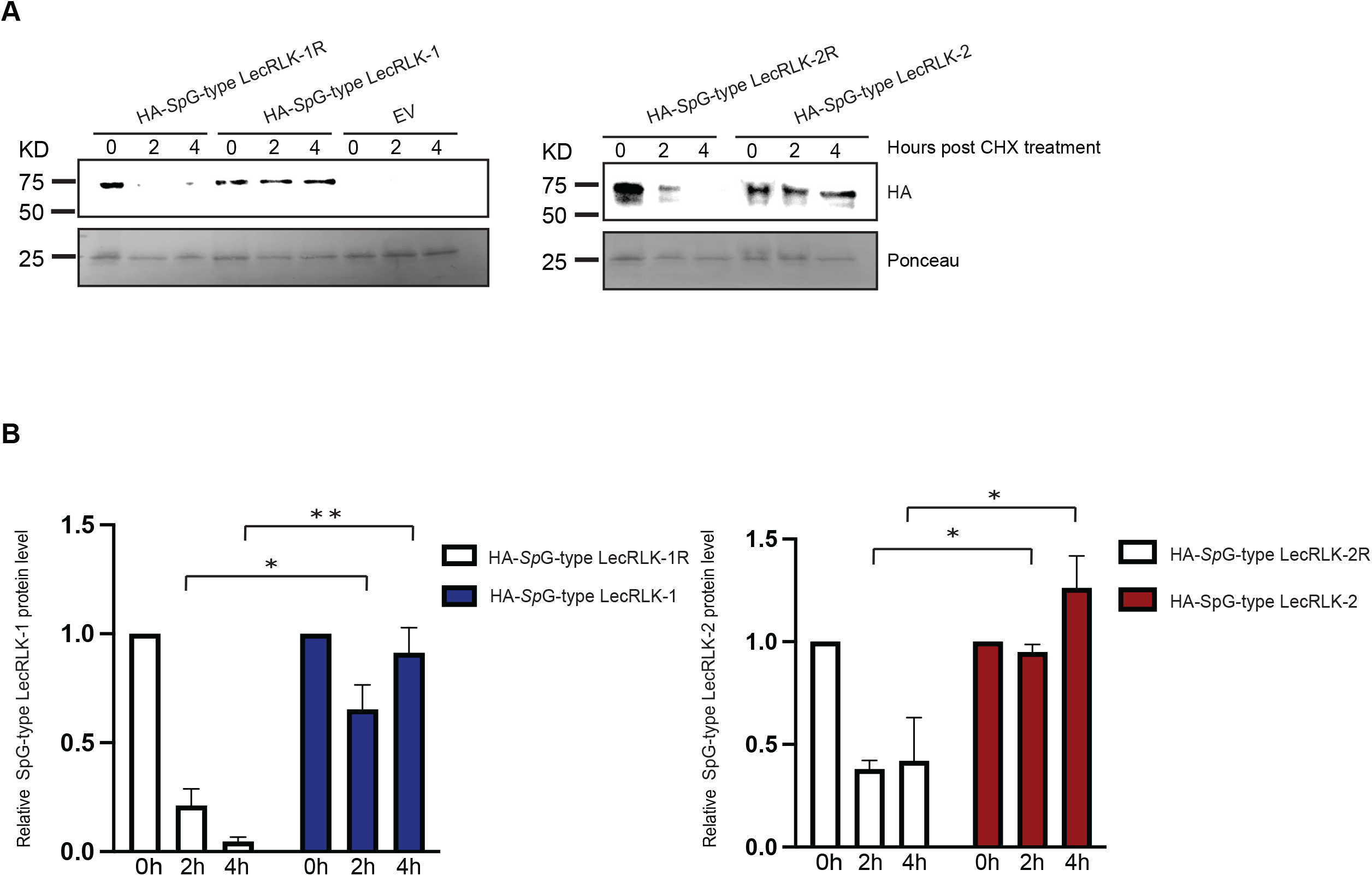
Conserved PAN domain residues modulates G-type Lec-RLK stability. A. 35S HA SpG-type LecRLK-1 and 35S HA SpG-type LecRLK-1R transformed Arabidopsis protoplast was pre-treated with cycloheximide and then the lysates were harvested as indicated time points. Total lysates were probed for HA. Ponceau staining used as internal control. Likewise, 35S HA SpG-type LecRLK-2 and 35S HA SpG-type LecRLK-2R transformed Arabidopsis protoplast were pre-treated with cycloheximide and then the lysates were harvested as indicated time points. Total lysates were probed for HA. Ponceau staining used as internal control. B. Band intensity were quantified using a densitometer and plotted relative intensities. Data are shown as mean ± SEM of three different independent experiments. Asterisks represents significant differences between transgenic mutant PAN and restored PAN plants (***P<0.0005; **P<.005; *P<0.05 One-way Anova, Sidak method).

The G-Lectin RLK family includes the S-locus receptor kinase (SRK), a receptor-like kinase which is involved in the self-incompatibility in the Brassica family (Sanabria, Goring et al. 2008). Previous studies confirmed that expressing recombinant SRK proteins in insect and *E.coli* cells lead to autophosphorylation on serine and threonine residues (Stein and Nasrallah 1993, Giranton, Dumas et al. 2000). These phosphorylation events were transphosphorylation, which suggested the existence of homo-oligomers of recombinant SRK proteins (Giranton, Dumas et al. 2000). Since we also confirmed higher ordered oligomers in the restored *Salix* G-type LecRLKs, we sought to investigate the phosphorylation status of the PAN mutated and PAN restored *Sp*G-type LecRLK-1. To determine the phosphorylation status of the variants, we overexpressed GFP-*Sp*G-type LecRLK-1 and GFP-*Sp*G-type LecRLK-1R in Arabidopsis. Western blot analysis revealed that the restored version had an increased phosphorylation level as determined by pan phosphoserine antibody (Figure 7A and 7B), suggesting that the PAN domain is essential for autophosphorylation of G-type LecRLKs.

**Figure 7:**
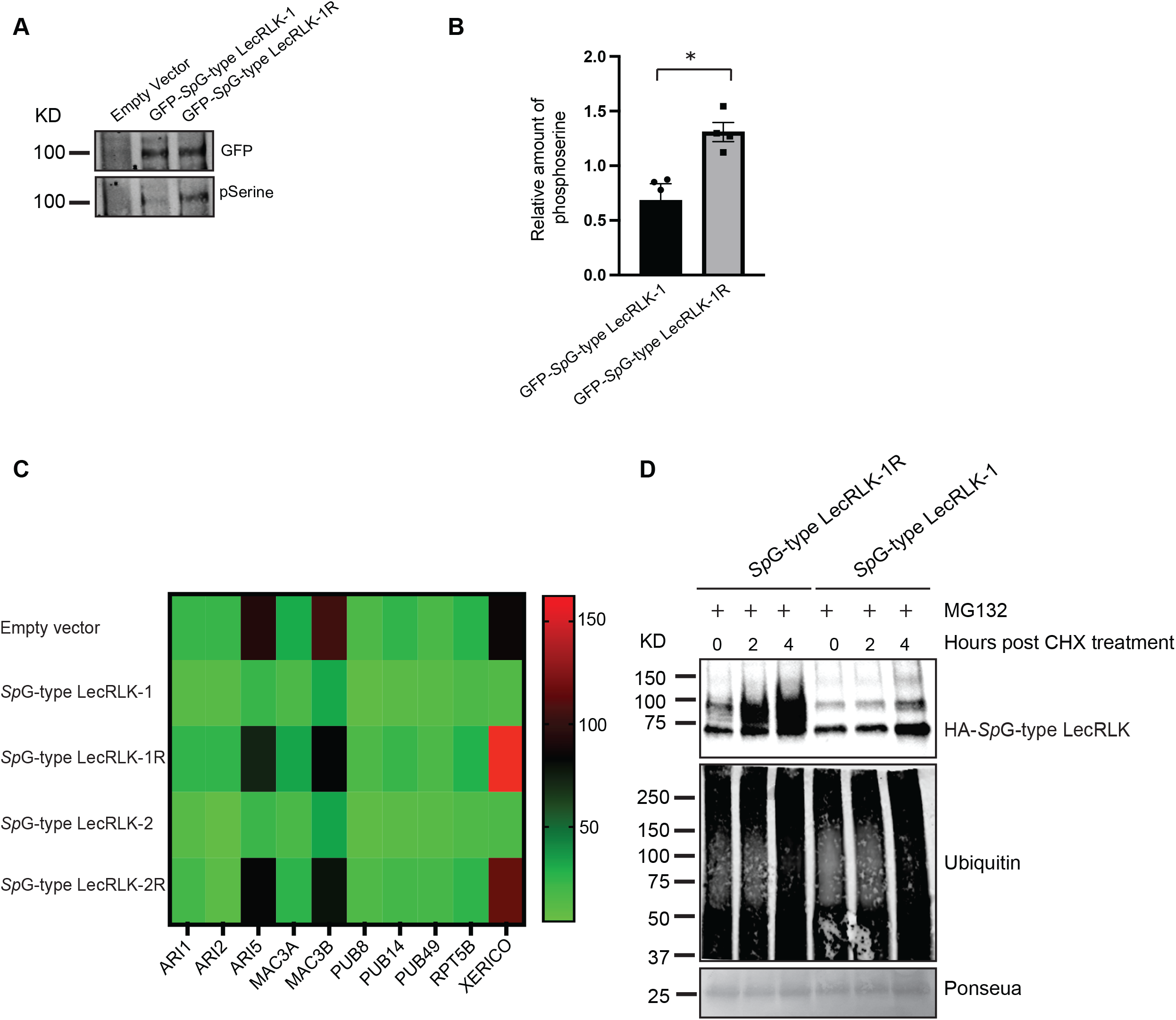
PAN domain is critical for G-type LECRLK phosphorylation and ubiquitination. A. GFP-tagged SpG-type LecRLK-1 and GFP-tagged SpG-type LecRLK-1R protein were detected with GFP antibody and probed subsequently with phosphoserine antibody. 3 biological replicates are shown. B. Band intensity were quantified using a densitometer and plotted relative intensities. Data are shown as mean ± SEM of three different independent experiments. Asterisks represents significant differences between transgenic mutant PAN containing G-type LECRLK and restored PAN containing G-type LecRLK plants (*P<0.05 Student T-test). C. Heat map of the differential expression analysis from RNA seq shows expression of E3 ubiquitin ligases in the empty vector, SpG-type LecRLK-1, SpG-type LecRLK-1R, SpG-type LecRLK-2 and SpG-type LecRLK-2R. D. 35S HA SpG-type LecRLK-1and 35S HA SpG-type LecRLK-1R transformed protoplast were treated with cycloheximide together with MG132 for 0, 2 and 4 hours. Ubiquitinated and total SpG-type LecRLK-1 and SpG-type LecRLK-1R proteins were detected with HA antibody on the lysate. The membrane was re-probed with ubiquitin specific antibody. Data are shown as representative of two different independent experiments.

We then assessed if there were any ubiquitin ligases that could be involved in the degradation pathway targeting the restored variants. Transcriptome analysis from Arabidopsis transgenic lines revealed several E3-ubiquitin ligases with higher expression levels in *Sp*G-type LecRLK-1R transgenic lines including ARI5, MAC3B and XERICO (Figure 7C). If indeed restored variants were being targeted for degradation, we hypothesized that MG132 which is well known proteasomal inhibitor could block the degradation of *Sp*G-type LecRLK-1R. As expected, pretreatment with MG132 resulted an accumulation of *Sp*G-type LecRLK-1R along with slower migration bands (Figure 7D). Since these slower migrating bands represents poly-ubiquitinated form of *Sp*G-type LecRLK-1R, these results suggest that an intact PAN domain in *Sp*G-type LecRLK-1R enhances homodimerization and leads to receptor ubiquitination prior to degradation by 26S proteasome, suggesting that the PAN domain is required for proteolytic degradation of *Sp*G-type LecRLK-1R.

### Conserved cysteine residues in PAN domain are critical for immunosuppression in plants

A characteristic pattern of the PAN domain is presence of 4-6 conserved cysteine residues that form its core (Figure 1). The PAN domain was implicated in protein ubiquitination and proteolysis of an oncoprotein, hepatocyte growth factor in human (Pal, De et al. 2022). Mutation of cysteine residues in the PAN domain of HGF led to major changes in HGF/cMET signaling in human cells. In addition to the HGF protein, mutating cysteine residues in a putative vestigial PAN domain in the human neuropilin protein, a receptor for SARS-Cov-2 binding and viral internalization, resulted is significantly reduced viral coat protein internalization (Pal, De et al. 2023). Based on these observations, we sought to evaluate the importance of conserved cysteine residues in the PAN domain of G-type LecRLK. To that end, we created 6 cysteine to 6 alanine mutation in both *Sp*G-type LecRLK-1R and *Sp*G-type LecRLK-2R variants (Figure 8A, 8C). Transgenic *Sp*G-type LecRLK-1PAN6cys-6Ala and *Sp*G-type LecRLK-2PAN6cys-6ala Arabidopsis plants showed induced transcript level of WRKY33 which suggests that conserved cysteines in the PAN domain of G-LecRLK also play a crucial role in immunosuppression (Figure 8B, 8D).

**Figure 8:**
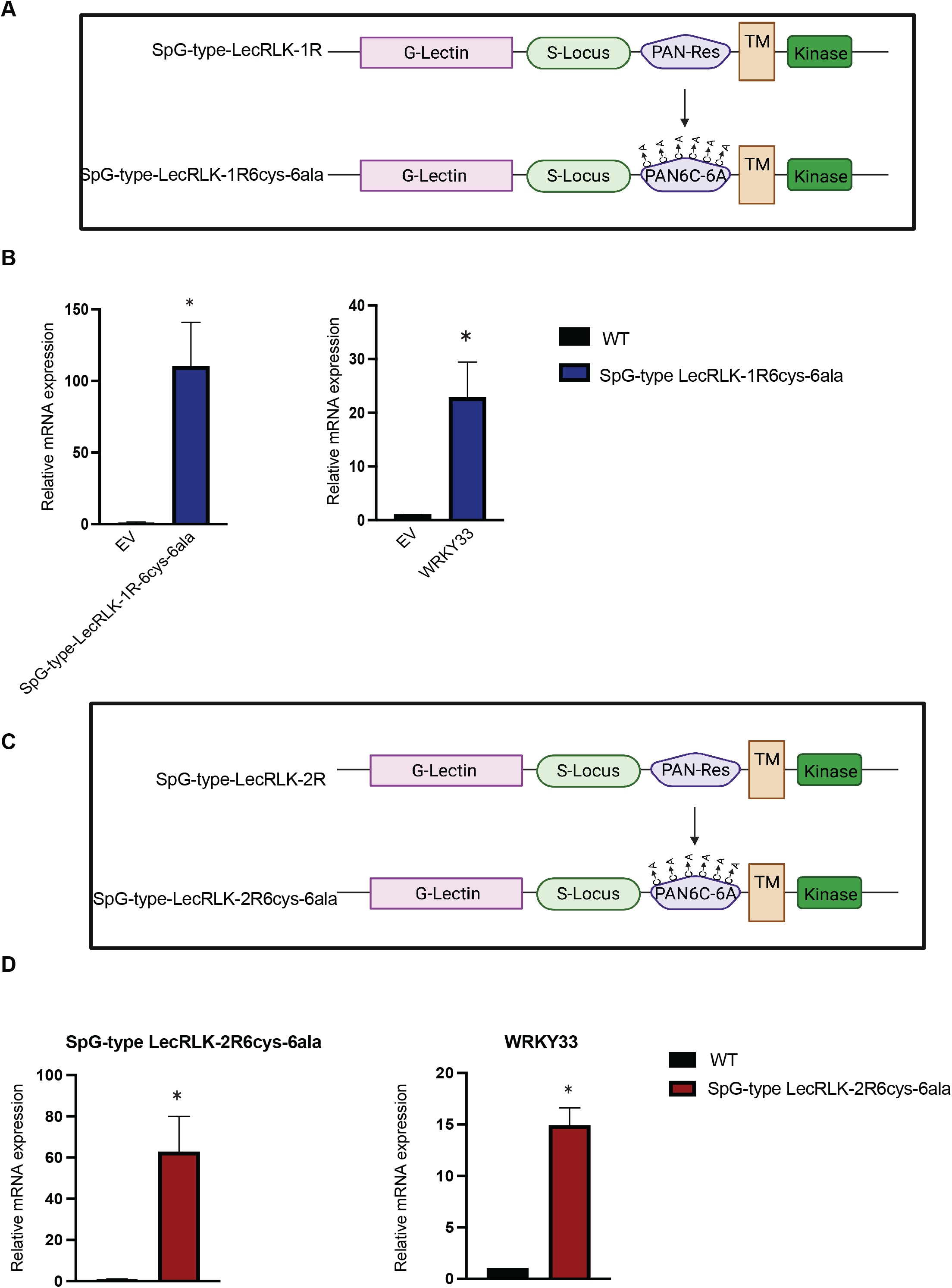
Conserved cysteine residues in PAN domain are critical for immunosuppression in plants. A. Schematic representation of transgenic SpG-type LecRLK-1R6cys-6ala construct from SpG-type LecRLK-1R. B. Quatitative real time PCR data from Arabidopsis transgenic plants shows induced expression of WRKY33 in transgenic SpG-type LecRLK-1R6cys-6ala Arabidopsis plants. Results are shown as mean ± SE of three different independent experiments. Asterisks represents significant differences between transgenic plants and wild type control (*P<0.05 Student T-test). C. Schematic representation of transgenic SpG-type LecRLK-2R6cys-6ala construct from SpG-type LecRLK-2R. D. Quatitative real time PCR data from Arabidopsis transgenic plants shows induced expression of WRKY33 in transgenic SpG-type LecRLK-2R6cys-6ala Arabidopsis plants. Results are shown as mean ± SE of three different independent experiments. Asterisks represents significant differences between transgenic plants and wild type control (*P<0.05 Student T-test).

### Homologous PAN domain present in *Arabidopsis* is conserved and exhibits similar mechanism in modulating plant immunity

After investigating the immune activation activities of overexpressed *SpG-type LecRLK1* and *SpG-type LecRLK-2* in Arabidopsis transgenic plants, we were interested to evaluate the function of PAN domain in *Arabidopsis* G-type LecRLK homolog. We took the advantage of the G-LecRLK, ERN1 which was recently shown as a negative regulator of Arabidopsis immunity against root-knot nematode *Meloidogyne incognita* also contains a PAN domain (Zhou, Godinez-Vidal et al. 2023). To perform this strategy, we aligned the *Sp*G-type LecRLK1 with the PAN domain of *Arabidopsis* ERN1 and At1g61380. After our domain prediction, we made minimal point mutations indicated by the arrows in the *ERN1* PAN domain (Figure S8A) to make it deactivated/mutated PAN domain. We used T-DNA insertions in ERN1, salk_128729 (*ern1.1*) which is essentially a null allele. To confirm the role of PAN domain, we generated transgenic complementation *Arabidopsis* lines, stably expressing ERN1 with functional PAN domain driven by ERN1 native promoter (ERN1pro::ERN1) and ERN1with mutated PAN domain driven by ERN1 native promoter (ERN1pro::ERN1 PAN mutated in knockout lines. We didn’t see significant difference between wild type Arabidopsis, *ern1.1* knockout lines, ERN1pro::ERN1, and ERN1pro::ERN1 PAN mutated lines (Figure S8E). However, ERN1pro::ERN1 lines show longer petioles compared to other variants. Real time PCR suggests ERN1pro::ERN1 PAN mutated lines displays triggered WRKY33 expression compared to wild type and ERN1pro::ERN1 variants (Figure 9A). Additionally, 35S pro::ERN1PAN mutated lines show increased WRKY33 expression compared to wild type and 35spro::ERN1 variants (Figure S8D). We were interested to measure the oxidative burst triggered by the perception of flg22 in a luminol dependent assay. This analysis showed that the ERN1pro::ERN1 PAN mutated transgenic lines had distinct enhancement of the oxidative burst compared to wildtype or transgenics expressing wild type version (Figure 9B). Additionally, we could see noticeable increase in pMAPK expression in ERN1pro::ERN1 PAN mutated transgenic lines compared to the wild type variant (FigureS8F). This result again bolsters our hypothesis that PAN domain modulates immune response in plants.

**Figure 9:**
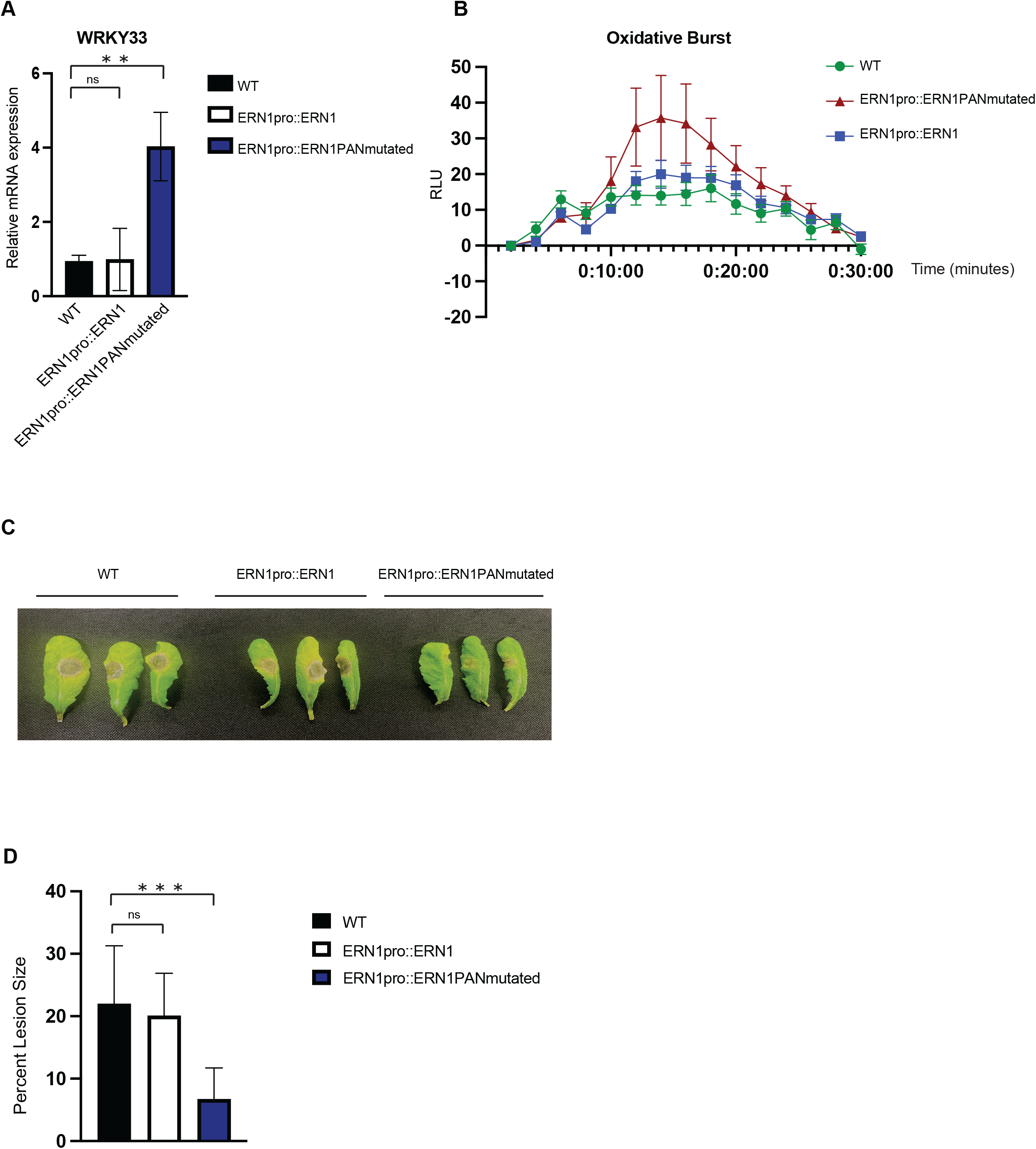
Expression analysis of transgenic ERN1pro: ERN1 and ERN1pro: ERN1 PAN mutated in *ern1.1* knock out *Arabidopsis*. A. Real time PCR for WRKY33 gene in ERN1pro: ERN1 and ERN1pro: ERN1 PAN mutated transgenic plants in *ern1.1* background. Asterisks represents significant differences between transgenic plants and wild type control (***P<0.0005; **P<.005; *P<0.05 Student T-test). B. Oxidative burst triggered by 500nM flg22 in WT, ERN1pro: ERN1 and ERN1pro: ERN1 PAN mutated in *Arabidopsis* transgenic plants measured in relative light units (RLU). Data represents standard deviation of the mean from three independent experiments from 9 biological replicates. C. Botrytis infection assay: Botrytis cinerea inoculation of WT, ERN1pro: ERN1 and ERN1pro: ERN1 PAN mutated in transgenic *ern1.1* Arabidopsis plants. D. Lesion area (percentage of total leaf area) in 72 hours post inoculation are represented in bar graphs. Value represents mean ± SE from at least n=20 from each group. Asterisks represents significant differences (***P<0.0005; **P<.005; *P<0.05 One-way Anova, Sidak method).

In summary, our study provides evidence that the PAN domain is required for suppression of JA and ET mediated defense signaling in plants. We presented molecular, genetic, transcriptomic, and biochemical evidence that naturally occurring mutations in conserved amino acid residues of the PAN domain in two *Salix* G-type LecRLKs trigger defense signaling when expressed in Arabidopsis and tobacco. Restoration to conserved amino acid residues disrupted defense signaling. We propose a model in which the PAN domain and its conserved amino acid residues promote suppression of JA and ET pathway by enhancing ubiquitination and proteolytic degradation of G-type LecRLKs to impair kinase activity and subsequent downstream signaling cascades (Figure 10A & 10B). Mutating the PAN domain retards receptor degradation (Figure 9B) and restores kinase activity to trigger downstream signaling cascades and JA and ET biosynthesis (Figure 9A). Based on our observations, we propose for the first time that the PAN domain mediates negative regulation of plant immunity by impairing JA and ET defense pathways.

**Figure 10:**
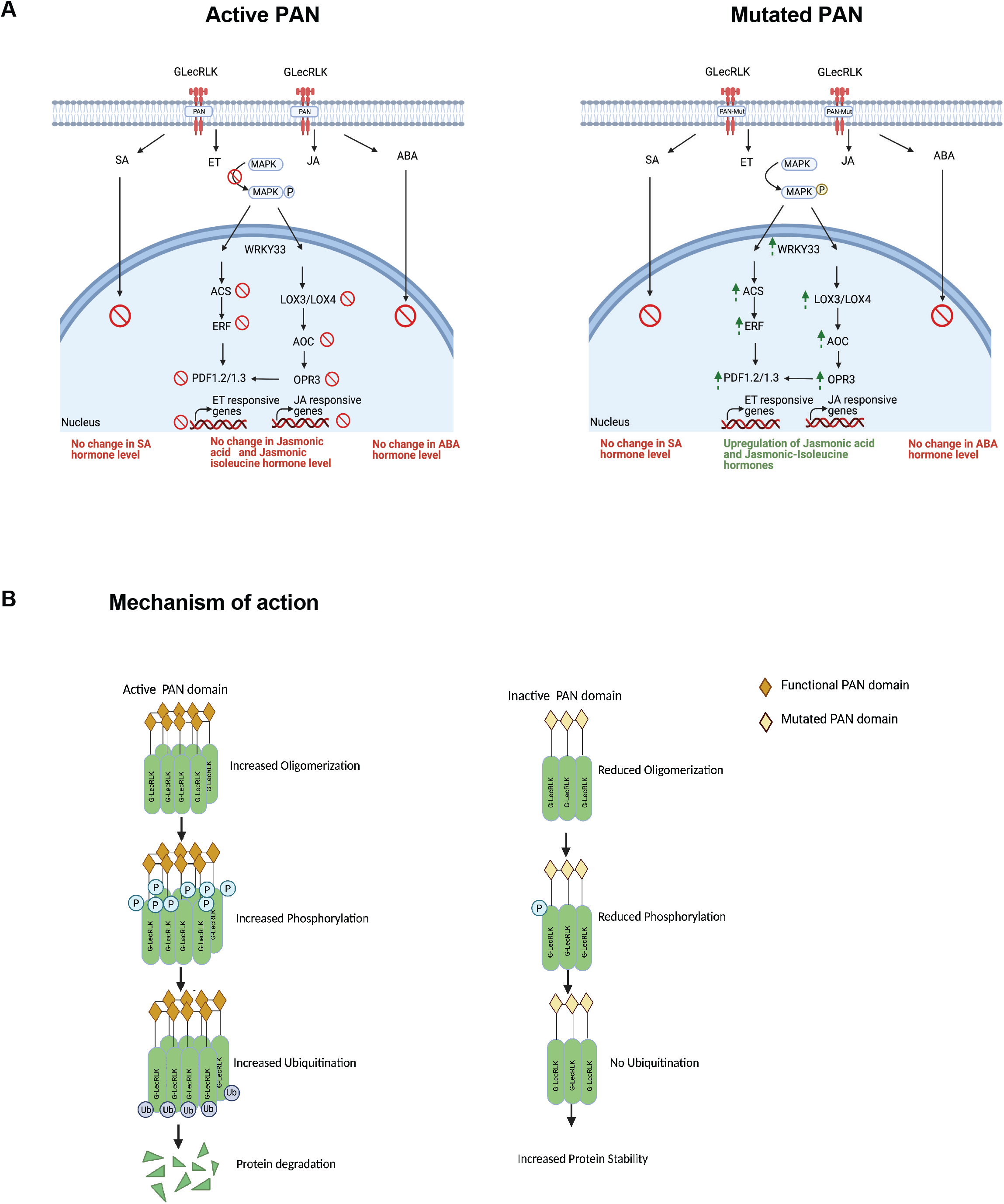
Working model to show the function of G-type LecRLK containing PAN domains in plants. A. Left: Schematic diagram indicates active PAN domain in transgenic SpG-type LecRLK-1 and SpG-type LecRLK-2 unable to activate JA and ET signaling, but mutation in conserved amino acid residues in PAN domain able to trigger the pathway (right). B. Working model showing the functional PAN domain of G-LecRLKs can self-interact and form higher order oligomers which gets targeted by ubiquitination and further suppresses plant immunity in host. On the other side, mutation in the conserved PAN domain of G-LecRLKs unable to form higher order oligomers and triggers host immunity in plants.

## Discussion

Multiple studies have shown that G-type LecRLK function as susceptibility factors for pathogens, symbionts and parasitic nematodes in plants. This phenomenon has not been widely studied and the exact mechanism behind it remains largely unknown. In this study, we demonstrated that the PAN domain plays an essential role in receptor turn-over to impair JA and ET signaling. Our observations that receptor ubiquitination and proteolysis occurred in the presence of an intact PAN domain whereas both processes were disrupted after mutating conserved amino acid residues in two different receptors offers a plausible mechanistic explanation. We propose that the PAN domain serves as direct or indirect target for ubiquitin ligases. This assertion is supported by transcriptome evidence which showed significant upregulation of multiple ubiquitin ligases when the PAN domain was restored. This led to markedly reduced protein abundance suggesting that both receptors were degraded before signaling cascades could be initiated. Retardation of signaling cascades was evidenced by lack of MAPK phosphorylation, WRKY33 expression remaining at wildtype levels and significantly less differentially expressed genes when G-type LecRLKs with restored PAN domains were overexpressed. In contrast, we observed robust MAPK phosphorylation, WRKY33 expression and global transcriptional reprogramming involving JA and ET defense genes when G-type LecRLKs with mutated PAN domains were overexpressed in Arabidopsis and tobacco. WRKY33 is a substrate of MPK3/MPK6 (Mao, Meng et al. 2011). Using a phospho-protein mobility shift assay, it has been demonstrated that WRKY33 is phosphorylated by MPK3/MPK6 in vivo in response to *Botrytis cinerea* infection (Mao, Meng et al. 2011). This study confirmed that WRKY33 functions downstream of MPK3/MPK6 in *Arabidopsis*. Previous study demonstrates that phosphorylation of WRKY33 is important for the activation of WRKY33 (Mao, Meng et al. 2011). Activation of WRKY33 specifically triggers genes involved in JA and ET pathway (Lorenzo, Piqueras et al. 2003) and is shown in (Figure 10A). Moreover, another study showed overexpression of ethylene response factor, ERF72 triggers camalexin biosynthesis and increased resistance to *B. cinerea*. The study specifically identified the transcription factor WRKY33 as a target genes which provides increased resistance to *B. cinerea* (Li, Liu et al. 2022). From these studies it shows that both JA and ET pathway signals through WRKY33. In our study, we demonstrate novel mechanism where the mutation of PAN domain in G-LecRLK phosphosphorylates MAPK3/6 which in turn specifically activates WRKY33 to activate JA and ET pathway.

Additionally, in this study, the function of PAN domain in Arabidopsis G-LecRLK receptor protein, ERN1 shows that mutation in the domain elevates immune response in plants against necrotrophic fungus *Botrytis cinerea.* Based on these observations, here, we propose that the PAN domain mediates negative regulation of plant immunity by impairing JA and ET defense pathways.

To date, the exact role of the PAN domain in protein function has remained vague with most studies only reporting that it is possibly involved in formation of disulfide bridges and could be involved in protein-protein or protein-carbohydrate interaction (Tordai, Bányai et al. 1999). A possible explanation as to why its function has remained elusive is that PAN domain-containing proteins appear to be largely unrelated and do not share clear evolutionary trajectories across divergent organisms. However, (Pal, De et al. 2021) revealed shared hallmark features of these proteins based on analysis of more than 28,000 PAN domain-containing proteins. These features included a predominant occurrence in the extracellular matrix and involvement in processes related to immune signaling. Notably, among the >28,000 is the Apical Membrane Antigen-1 (AMA1) protein which is found in apicomplexan parasites including *Toxoplasma gondii* and *Plasmodium spp*. AMA1 has been widely shown to be essential for attachment and host-cell invasion by these parasites (Tyler, Treeck et al. 2011), a process that is akin to G-type LecRLK-mediated infection of plants by microbes. Another example involves PAN domain-containing zona pellucida glycoproteins which surround mammalian female eggs and function as sperm receptors during gamete fertilization (Gupta 2015). Taken together, these cumulative observations suggest that the PAN domain might serve the same conserved function in facilitating host-cell invasion across divergent biological systems.

## Method

### Plant material and growth condition

*Arabidopsis thaliana*, Columbia ecotype and *Nicotiana benthamiana* plants were grown in Metro-Mix 200 soil or on germination plates (Murashige and Skoog) a growth chamber at 22°C with a 16-h-light/8-h-dark cycle. Leaves were collected from rosette-stage plants grown on soil and used for DNA isolation, and genotyping. For transcript and protein analysis all samples were harvested, frozen in liquid N_2_, and stored at −80°C until needed. The coding sequences of *Sp*G-type LecRLK-1 (SapurV1A.0918s0020), *Sp*G-type LecRLK-2 (SapurV1A.003s0270), and SapurV1A.1175s0020 were amplified by PCR from cDNA derived from *Salix purpurea* leaves using HiFi polymerase (Clontech). Sequences were determined using Phytozyme using the Salix purpurea v.1.0 genome. The pENTR/D-TOPO entry vector was used for the insertion of the PCR products using the pENTR Directional Topo Cloning Kit (Thermo Fisher Scientific). LR Clonase II Enzyme Mix (Thermo Fisher Scientific) was used to transfer the coding sequences into the pGWB402-omega destination vector and transformed into One Shot TOP10 Chemically Competent cells. Vectors were transformed into the *Agrobacterium* strain GV3101 and transformed into Col-0 Arabidopsis thaliana using the floral dip method. Transformed seeds were selected with Kanamycin and grown in long day growth chambers at 16/8 hours light/dark at 22°C. Likewise restored versions, *Sp*G-type LecRLK-1R and *Sp*G-type LecRLK-2R cloned into binary vector pGWB402 omega for transformation. *ERN1* CDS and *ERN1* CDS with deactivated PAN domain was cloned in pGWb402Omega under 35S promoter. Full length *ERN1* and *ERN1* with mutated PAN domain with 1,168bp upstream promoter fragment was cloned in pGWb540 under native promoter. The complemented transgenic lines were screened on hygromycin containing Murashige and Skoog medium.

All the primers used to make these constructs are present in TableS1.

### Pathogen infection and analysis

Four-week-old Arabidopsis leaves were detached and inoculated with 5 µl *Botrytis cinerea* conidiospore suspension (5 × 10^5^ spores ml^−1^ in potato dextrose broth) as previously described to analyze the disease symptoms (Ingle and Roden 2014). Leaf lesions were pictured and measured using Image J software. For each genotype, at least 20 independent values were used for statistical analysis.

### Protoplast isolation and transfection

Arabidopsis mesophyll protoplasts were isolated and transfected as described previously (Yoo, Cho et al. 2007). Protoplasts were isolated from fully expanded leaves from 3–4-wk-old Col-0 plants. For Cycloheximide and MG132 assay 20 µg of *SpG-type LecRLK* DNA were transfected into 200 µl of protoplasts. After 18 h incubation, protoplasts were treated with MG-132 and Cycloheximide as indicated and were collected accordingly. Total proteins were extracted using protein extraction buffer for western blot analysis.

### RNA Extraction and qRT-PCR

Three-week-old leaf tissue was collected for RNA extractions using the Spectrum Plant Total RNA Kit (Sigma) and cDNA synthesized using the Superscript III Reverse Transcriptase with oligo-d(T) primers (Thermo Fisher Scientific). qRT-PCR was performed using the PowerUP SYBR Green Master Mix (Thermo Fisher Scientific) and relative expression for each gene was determined using 2ΔΔCt method with GAPDH as an internal control. All the primers used in the study is present in table S1.

### Phytohormone Analysis

*Arabidopsis thaliana* using electrospray ionization-high-performance liquid chromatography tandem mass spectrometry (ESI-HPLC-MS/MS). The plant hormones include jasmonic acid (JA), jasmonoyl-L-isoleucine (JA-ILE), abscisic acid (ABA), and salicylic acid (SA).

A. Preparation of Standard Solution of Plant Hormones

1) 2 μL of 500 μg/ mL standard reserve solution for each hormone was added to 986 μL methanol in a 1.5 mL centrifuge tube. The solution was mixed well and prepared as the stock solution with a final concentration of 1 μg/ mL. 2 μL of 500 μg/ mL internal standard reserve solution for each hormone was added to 996 μL methanol in a 1.5 mL centrifuge tube. The solution was mixed well and prepared as the stock solution with a final concentration of 1 μg/ mL. Then, 989.9 μL, 989.8 μL, 989.5 μL, 988 μL, 985 μL, 970 μL, 940 μL, 790 μL methanol was added into 1.5 mL centrifuge tubes, and the stock solution prepared in step (1) was added into the methanol solution in sequence of 0.1 μL, 0.2 μL, 0.5 μL, 2 μL, 5 μL, 20 μL, 50 μL, 200 μL, respectively. 10 μL of the internal standard in step (2) was added to each tube to make standard solution with final concentration 0.1 ng/ mL, 0.2 ng/mL, 0.5 ng/mL, 2 ng/mL, 5 ng/mL, 20 ng/mL, 50 ng/mL, 200 ng/mL with 10 ng/mL internal standard.

### Steps for Phytohormones Extraction

1. About 1 g sample was weighed, grinded, and added into 10 times volume of acetonitrile solution with 4 μL internal standard stock solution.
2. The sample solution was briefly vortexed and kept at 4 °C overnight.
3. Following centrifugation for 5 min at 12,000 g at 4 °C, the precipitate was extracted once again using 5 times volume of acetonitrile solution.
4. 35 mg C18 packing was added and mixed vigorously for 30 s, centrifuged at 10,000 g for 5 min, and the supernatant was saved.
5. The supernatant was dried by nitrogen and then dissolved in 400 μL methanol.
6. After filtration through a membrane filter (0.22 μm), 2 μL extract was injected onto a C18 column mounted on an analytical HPLC system (AGLIENT) equipped with AB SCIEX-6500 Qtrap MS/MS.

### Ethylene measurement Assay

Seedlings were grown on agar in jelly jars with a 390 mL headspace. 21 day old seedlings were sealed with a lid fitted with a septum and ethylene allowed to accumulate for 60 min at which time the concentration of ethylene in each jar was determined with an ETD-300 laser detector (Sensor Sense, The Netherlands).

### Western blotting and immunoprecipitation

FLAG tagged *Sp*G-type LecRLK-1 and *Sp*G-type LecRLK-1R and GFP-tagged *Sp*G-type LecRLK-1 and *Sp*G-type LecRLK-1R were expressed where indicated in 293T cells. Cell extracts were generated in EBC buffer, 50 mM Tris (pH 8.0), 120 mM NaCl, 0.5% NP40, 1 mM DTT, and protease and phosphatase inhibitors tablets (Thermo Fisher Scientific). For protein phosphorylation, GFP tagged SpG-type LecRLK-1, and SpG-type LecRLK-1R were expressed in Arabidopsis. Protein extracts were generated using Sigma plant protein extraction kit (PE0230) from the transgenic plants. The isolated proteins were probed with GFP and pSerine antibody subsequently. For immunoprecipitation, equal amounts of cell lysates were incubated with the indicated antibodies conjugated to protein G beads (Invitrogen) (15μl per IP, Thermo Scientific) overnight at 4 °C. The beads were then washed with EBC buffer including inhibitors. Immunoprecipitation samples or equal amount of whole-cell lysates were resolved by SDS-PAGE, transferred to PVDF membranes (Millipore) probed with the indicated antibodies, and visualized with the LiCor Odyssey infrared imaging system.

### Mammalian cell culture and transfection

HEK 293T cells were obtained from ATCC and maintained in a humidified atmosphere at 5% CO2 in Dulbecco’s Modified Eagle’s (DMEM) complete medium (Corning) supplemented with 10% fetal bovine serum (FBS; Seradigm) in 37°C. Plasmid transfections were done with TransIT-LT1 (Mirus Bio) per the manufacturer’s instructions.

### Antibodies

HA (#902302; 1:1000) antibody was purchased from Biolegend. FLAG antibody (F3165) from Sigma, phosphoserine antibody from Sigma Aldrich clone A4 (05-1000X), and GFP antibody (A11122) from Invitrogen. p44/42 MAPK (Erk1/2) (#9102; 1:1000), Ubiquitin (E412J) mAb #43124 were purchased from Cell signaling. Chemiluminescence detection was performed according to the manufacturer’s instructions (Amersham ECL Western Blotting Detection Reagent kit) followed by exposure using Chemidoc gel-documentation system (BioRad). For imaging using Li-Cor, secondary antibodies for western blotting were purchased from LI-COR Biosciences.

### Cloning, heterologous protein expression, purification, and size exclusion chromatography

The PAN-like domain of *Sp*G-type LecRLK-1 and *Sp*G-type LecRLK-1R (amino acids 340-448) were amplified from plasmid templates pGWb402Omega-*Sp*G-type LecRLK-1 and pGWb402Omega-*Sp*G-type LecRLK-1R, respectively, using the following primers: AA340F 5’-AACTTGTACTTTCAAGGCTACCAGGATTCGGTAAAT-3’ and RLK7_AA448R 5’-ACAAGAAAGCTGGGTCCTACACTTTTAAGAATGAAGT −3’. Likewise, the PAN domain of *Sp*G-type LecRLK-2 and *Sp*G-type LecRLK-2R (amino acids 322-400) was amplified from plasmid templates pGWb402Omega-*Sp*G-type LecRLK-2 and pGWb402Omega-*Sp*G-type LecRLK-2R, respectively, using the following primers: RLK5_AA322F 5’-AACTTGTACTTTCAAGGCGCTCTGAATTGTGATTCC-3’ and RLK5_AA400R 5’-ACAAGAAAGCTGGGTCCTAAGATATCGAGCTTAAAGA-3’ To create Gateway entry clones, *att*B-PCR products were generated using two-step adapter PCR (Prabhakar, Wang et al. 2020). The resulting expression constructs encode fusion proteins comprised of an NH_2_-terminal signal sequence, an 8xHis tag, an AviTag recognition site, the “superfolder” GFP (sfGFP) coding region, the recognition sequence of the tobacco etch virus (TEV) protease, followed by residues 322-400 of *Sp*G-type LecRLK-2 (or the *Sp*G-type LecRLK-2R variant) or residues 340-448 of *Sp*G-type LecRLK-1 (or the *Sp*G-type LecRLK-1R variant). Recombinant expression was accomplished by transient transfection of HEK293 cells (FreeStyle 293-F cells, ThermoFisher) at a 1L scale. Soluble secreted fusion proteins were purified from the culture media using HisTrap HP columns (Cytiva, USA) with an ÄKTA Pure 25L protein purification system (Cytiva, USA) as described previously (Prabhakar et al., 2020), and quantified using GFP fluorescence. The eluted fractions (more than 90% pure), identified by measuring GFP fluorescence. For high-resolution preparative gel filtration chromatography, 5 mL of purified, concentrated protein was injected onto a HiLoad 16/600 GL, Superdex 75 pg column (Cytiva, USA) pre-equilibrated with 50 mM HEPES, 400 mM NaCl using an ÄKTA Pure 25L protein purification system (Cytiva, USA). 5 mL fractions were collected based on absorbance at 280 nm with 5 mAU as a cutoff. All protein purification steps were carried out at 4°C.

### RNA-Seq Experiment

RNA was extracted from transgenic tobacco and Arabidopsis leaves using the Sigma plant RNA extraction kit. Quantification and quality control of isolated RNA was performed by measuring absorbance at 260 nm and 280 nm on a NANODROP ONEC spectrophotometer (Thermo Scientific, USA). mRNA molecules were purified from total RNA using oligo(dT)-attached magnetic beads. Following that, mRNA molecules were fragmented into small pieces using fragmentation reagent after reaction a certain period in proper temperature. First-strand cDNA was generated using random hexamer-primed reverse transcription, followed by a second-strand cDNA synthesis. The synthesized cDNA was subjected to end-repair and then was 3’ adenylated. Adapters were ligated to the ends of these 3’ adenylated cDNA fragments. This process was to amplify the cDNA fragments with adapters from previous step. PCR products were purified with Ampure XP Beads (AGENCOURT) and dissolved in EB solution. Library was validated on the Agilent Technologies 2100 bioanalyzer. The double stranded PCR products were heat denatured and circularized by the splint oligo sequence. The single strand circle DNA (ssCir DNA) were formatted as the final library. The library was amplified with phi29 to make DNA nanoball (DNB). The DNBs were load into the patterned nanoarray and 150 paired-end reads were generated in the way of sequenced by synthesis. The RNA-seq run was performed with four biological replicates. The raw RNA-seq reads have been deposited at NCBI under BioProject ID PRJNA739633. Clean RNA-Seq reads were mapped to the TAIR10 *A. thaliana* reference genome with bowtie2 v2.2.5(Langmead and Salzberg 2012). Expression level as Fragments per kilobase per million reads (FPKM) were generated with RSEM v1.2.8 (Li and Dewey 2011). Differential expression analysis was performed with DESeq2, genes with an adjusted p-value less than 0.05 were considered differentially expressed (Love, Huber et al. 2014).

### PAN domain alignment

PAN domains in *Sp*G-type LecRLK-1 (SapurV1A.0918s0020), *Sp*G-type LecRLK-2 (SapurV1A.0037s0270), *Potri*.T022200, *Sp*G-typeV1A.1175s0020 (SapurV1A.1175s0020), *Sp*G-typeV1A.1316s0010 (SapurV1A.1316s0010), and AT1G61380 were identified with InterProScan 5(Jones, Binns et al. 2014). The PAN domain region was extracted from the full-length protein and aligned with MAFFT *linsi(Katoh and Standley 2013)*. The alignment was visualized with Geneious Prime (Kearse, Moir et al. 2012). PAN domain of *Sp*G-type LecRLK-1, At1g61380 and ERN1 was aligned using PRALINE multiple sequence alignment. The level of conservation determined by different color mode.

### Measurement of ROS Generation

Reactive oxygen species released by leaf tissue was determined by hydrogen peroxide-dependent luminescence of luminol. Leaf discs of 5-mm diameter from 4-week-old plants were punched and floated overnight in darkness in 96-well plates on 100ul distilled water. Horseradish peroxidase (10 µg/mL) and luminol (100 µm) from Sigma-Aldrich was added for elicitation and ROS .detection. Luminescence was measured directly after addition of 500uM elicitor peptide Flg22 in plate reader Biotek Synergie2 for specified timepoints. For the H_2_DCFDA experiment, protocol was adapted from a previous report(Leshem, Melamed-Book et al. 2006). Briefly, 10-d-old seedlings, acclimated for 48 h in liquid MS medium, were incubated in 10 μm H_2_DCFDA(Sigma-Aldrich) in phosphate-buffered saline solution for 30 min and washed twice for 5 min in phosphate-buffered saline solution. Chlorophyll autofluorescence and oxidized H_2_DCFDA were visualized using a Zeiss confocal microscope. H_2_DCFDA fluorescence was detected at 535 nm and chlorophyll at 650 nm and over.

### Statistical Analysis

Statistical analyses were performed on individual experiments, as indicated, with the GraphPad Prism Software using an unpaired t-test. Sample sizes and specific tests are indicated in the figure legends. A p-value of <0.05 was considered significant.

## Acknowledgements

This research was supported by the United States Department of Energy’s Office of Science Early Career Research Program under the Biological and Environmental Research office and National Institutes of Health (P41GM103390 and R01GM130915). Oak Ridge National Laboratory is managed by UT-Battelle, LLC for the U.S. Department of Energy under Contract Number DE-AC05-00OR22725. Part of this work was performed at the Oak Ridge Leadership Computing Facility (OLCF). We thank Dr. Lawrence Smart at Cornell University for providing *Salix* propagules. We thank Chandler Brown for maintaining *Arabidopsis* transgenics and Dr. Christopher A. Makaroff at Miami University for critical comments and editing of the manuscript. We made schematic models using Biorender.com.

## Author Contributions

WM supervised the whole study and all the experiments on the paper. KD and WM performed analysis of all the figures. KD, CS, TY and KF contributed to bioinformatics analysis and KD and TY performed RNA seq analysis; KD, DP and SJ contributed to isolate RNA; KD and DP performed the biochemical assays; KD and DP performed the proteasomal degradation assay; KD contributed to physiological analysis; KD performed the pathogenic response assay; KD and DP contributed to imaging and imaging analysis; KD and BB performed the ethylene assay; KD performed phytohormone analysis; KD, CS, contributed to generate transgenics in Arabidopsis; KD contributed to generate transgenics in tobacco and KD and SJ contributed in tobacco maintenance; KD, DP, CS, MH, generated clones for the study; DP and KD expressed the constructs in mammalian cells for co-immunoprecipitation; JYY, DK and KWM expressed the recombinant products in mammalian cells, PKP and BU purified the recombinant products and contributed to the size exclusion chromatography experiments; KD and DP made all the figures; KD, DP and WM drafted the manuscript and all authors critically reviewed and approved the final version of the manuscript for publication.

All Revision Work: KD and DP, supervised by WM.

